# eDITH: an R-package to spatially project eDNA-based biodiversity across river networks with minimal prior information

**DOI:** 10.1101/2024.01.16.575835

**Authors:** Luca Carraro, Florian Altermatt

## Abstract

1. Ecological and ecosystem monitoring is rapidly shifting towards using environmental DNA (eDNA) data, particularly in aquatic systems. This approach enables a combined coverage of biodiversity across all major organismal groups and the assessment of ecological indices. Yet, most current approaches are not exploiting the full potential of eDNA data, largely interpreting results in a localized perspective. In riverine networks, by explicitly modelling hydrological transport and associated DNA decay, hydrology-based models enable upscaling eDNA-based diversity information, providing spatially integrated inference. To capitalize from these unprecedented biodiversity data and translate into space-filling biodiversity projections, a streamlined implementation is needed.
2. Here, we introduce the `eDITH` R-package, implementing the eDITH model to project biodiversity across riverine networks with minimal prior information. eDITH couples a species distribution model relating a local taxon’s eDNA shedding rate in streamwater to environmental covariates, a mass balance expressing the eDNA concentration at a river’s cross-section as a weighted sum of upstream contributions, and an observational model accounting for uncertainties in eDNA measurements. By leveraging on spatially replicated eDNA measurements and minimal hydromorphological data, eDITH enables disentangling the various upstream eDNA sources, and produces space-filling maps of a taxon’s spatial distribution at any chosen resolution. eDITH is applicable to both eDNA concentration and metabarcoding data, and to any taxon whose DNA can be retrieved in streamwater.
3. The `eDITH` package provides user-friendly functions for single-run execution and fitting of eDITH to eDNA data with both Bayesian methods (via the `BayesianTools` package) and non-linear optimization. An interface to the `DHARMa` package allows model validation via posterior predictive checks. Necessary preliminary steps such as watershed delineation and hydrological characterization are implemented via the `rivnet` package. We illustrate `eDITH`’s workflow and functionalities with two case studies from published fish eDNA data.
4. The `eDITH` package provides a user-friendly implementation of eDITH, specifically intended for ecologists and conservation biologists. It can be used without previous modelling knowledge but also allows customization for experienced users. Ultimately, eDITH allows upscaling eDNA biodiversity data for any river globally, transforming how state and change of biodiversity in riverine systems can be tracked at high resolution in a highly versatile manner.

## 1 Introduction

Environmental DNA (eDNA, that is, DNA isolated from environmental samples; Taberlet et al. 2012; Pawlowski et al. 2020) has revolutionized ecological research, allowing for fast, cost-effective and taxonomically broad biodiversity assessments, especially in aquatic media (Thomsen and Willerslev, 2015; Valentini et al., 2016; Deiner et al., 2017; Beng and Corlett, 2020). Notably, in lotic environments, owing to dynamical hydrological processes, eDNA aggregates biodiversity information across broad contributing areas (Deiner and Altermatt, 2014; Barnes and Turner, 2015; Deiner et al., 2016; Shogren et al., 2017; Seymour et al., 2021). As such, eDNA has advanced our understanding of biodiversity structure and ensuing ecosystem functions in river ecosystems (Altermatt et al., 2020). Its game-changing potential is directly linked to the passive transport of eDNA in streamwater, providing a spatial integration of biodiversity information. This approach is paralleling the revolution in environmental chemistry, where local samples can be used to spatial infer sources and contamination levels (Abbott et al., 2018; Cairoli et al., 2023; Fairbairn et al., 2016). In analogy, the occurrence of single organisms, whole communities and derived ecological indices can be attributed and spatially assigned in any riverine networks.

Such an attribution has been implemented for the first time by the development of the eDITH model (*eD*NA *I*ntegrating *T*ransport and *H*ydrology; Carraro et al. 2017, 2018, 2020b, 2021, 2023), which couples a geomorphological and hydrological characterization of a watershed with a species distribution model and eDNA transport and decay dynamics. This model uniquely leverages spatially replicated eDNA measurements within a river network to infer the spatial distribution of any taxon of interest, expressed as relative density and/or detection probability. The eDITH model can be applied to any taxonomic unit (hereafter “taxon”), including Operational Taxonomic Units (OTUs) or Amplicon Sequence Variants (ASVs), and can handle DNA concentration data (e.g., from quantitative Polymerase Chain Reaction qPCR) and metabarcoding (read counts) data. Hitherto, this integration has been only accessible to specialist users with extensive combined hydrological and ecological knowledge, and depended on customized watershed delineation, hydrological characterization, model implementation and fitting.

Here, we present `eDITH`, an R-package for simple and direct application of eDITH, allowing a direct workflow for translating eDNA point data into space-filling biodiversity maps across watersheds based on minimal input information. The package exploits a seamless R-based river network extraction provided by the `rivnet` package (Carraro, 2023), only requiring the coordinates of two points defining the spatial range of the watershed of interest and one point identifying the watershed outlet. Hydraulic variables necessary to the eDITH model are extrapolated based on a single measure of stream width and discharge. As such, `eDITH` allows a user-friendly implementation of eDITH, thus making it accessible to a wide array of end users without depending on extensive knowledge of hydrology, ecological modelling or molecular ecology. For the more experienced users, the package enables detailed customization, for instance in terms of choices of model fitting technique, covariates and error distribution. In the following, we outline the model’s underlying assumptions and applicability, as well as its mathematical foundation; we then describe the package structure and its operational workflow, and conclude by briefly illustrating two case studies.

## 2 Model overview

### 2.1 Underlying assumptions and applicability

The key underlying concept of eDITH is that DNA particles are advected downstream by streamflow. Consequently, an eDNA sample is not only representative of the location where it is taken, but it provides information about taxon abundance (in the case of single-species data obtained via e.g. qPCR) or biodiversity (for metabarcoding data) for a certain area upstream of the sampling location. Advection of eDNA with streamflow has been extensively documented (Barnes and Turner, 2015; Civade et al., 2016; Deiner and Altermatt, 2014; Deiner et al., 2016; Pont et al., 2018; Sansom and Sassoubre, 2017; Shogren et al., 2017; Zhang et al., 2023), yet has only been minimally exploited to infer information on eDNA sources. By exploiting information from multiple sampling sites distributed in space, and embedding a model for transport and concurrent decay of DNA, eDITH disentangles the various sources of DNA shedding, and hence, the spatial distribution of the target taxon’s abundance. As a result, eDITH complements point-wise eDNA measurements by spatially projecting taxon distributions (and thus biodiversity information) into space-filling catchment maps. This approach is a twofold capitalization of eDNA-based diversity assessment compared to any traditional monitoring. Firstly, it is generalizable across any taxonomic group. Secondly, it upscales the understanding of species occurrences and derived biodiversity or ecological indices from a point-based distribution to a space-filling projection.

#### DNA production and decay rates

The DNA production (i.e., shedding) rate of a taxon in streamwater is assumed to be proportional to the taxon’s density (Lodge et al., 2012; Apothéloz-Perret-Gentil et al., 2017), which can be equivalent to both biomass and abundance. While shedding rates scale allometrically with organismal mass (Yates et al., 2021), the distinction between biomass and abundance is, for modelling purposes, irrelevant. In essence, the patterns modelled by eDITH are those of a quantity that is proportional to the DNA production rate–be it biomass, abundance or allometrically scaled mass. The validity of this assumption is restricted to the time point when eDNA is sampled. Thus, it is irrelevant if a taxon varies in its DNA release rate across a season, as long as its released DNA is proportional to its density at the time of sampling. For the same reason, hydrological variables, such as discharge and velocity, are assumed constant in time (but not in space) across the duration of a sampling campaign. For instance, if multiple days are required to perform eDNA sampling across all sampling sites, this period would be considered as a unique time point, during which hydrological conditions are adequately represented by time-averaged discharge and velocity values, thus implying that sampling should be done at constant, base-flow conditions. The decay of DNA is assumed to follow first order kinetics, i.e., the rate of change of DNA concentration decreases linearly with time. Similarly to the production rate, a single value of decay time is assumed as representative for a given taxon at a given time point.

#### Sampling design

The spatial extent of eDNA sampling must adequately cover the entire river network to allow robust model predictions. As a rule of thumb, there should be one sampling site per each 10– 20 km^2^ of drained area. Moreover, all main tributaries of a river network should be sampled in a spatially hierarchical and nested design. Ideally, sampling sites should be located just upstream of a confluence, so that the independent signals from the joining tributaries can be gauged independently (Fig. 1). Details on optimal eDNA sampling strategies are discussed in Carraro et al. (2021).

**Figure 1:**
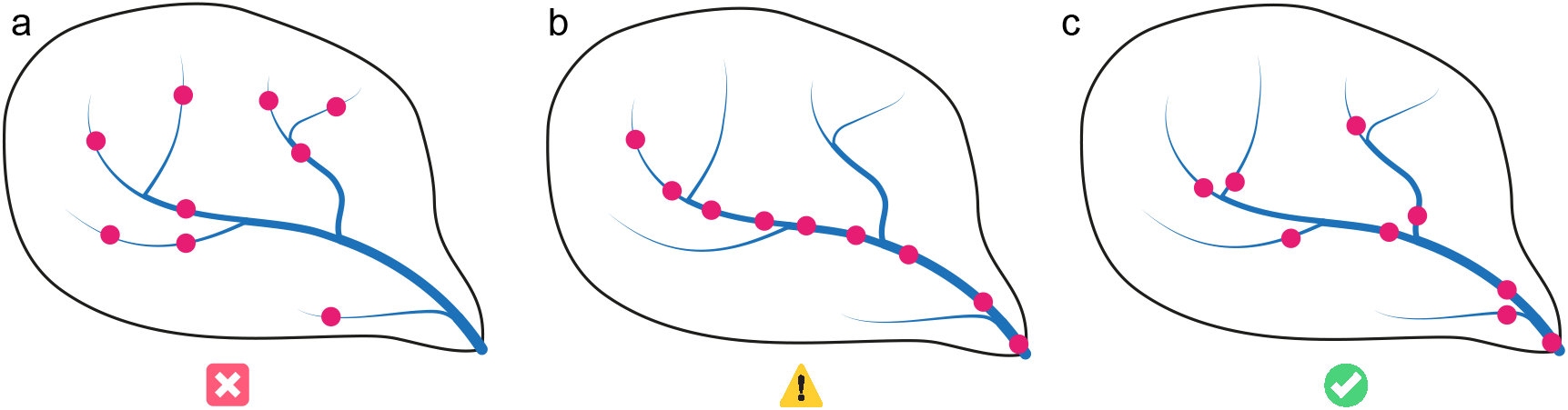
Comparison of sampling designs for eDNA collection in river networks. a) An upstream-heavy sampling strategy, with limited nestedness (i.e., the degree to which sampling sites are connected by downstream paths) is poorly suitable for application of eDITH. b) A strategy with samples taken only along the main stem, and thus high nestedness. This allows eDITH to provide robust estimates of the taxon distribution along the main stem, but the prediction power for other tributaries will be limited. c) A more downstream-heavy strategy, with sites located at the downstream end of the respective reach (i.e., right upstream of a confluence). This is the best sampling strategy, as it allows eDITH to disentangle the various eDNA sources across the river network.

The mean along-stream distance between eDNA samples must be short enough that the eDNA found at an upstream site is not totally depleted before reaching the next downstream site (Deiner and Altermatt, 2014; Pont et al., 2018). Thus, transport and decay must allow a spatially dependent signal between individual sampling sites, otherwise eDITH can not make inferences based on multiple sites. As an example, under first order kinetics, the eDNA concentration *C*_*L*_ at a distance *L* downstream of a reference point is *C*_*L*_ = *C*_0_ exp [−*L*/(*vτ*)], where *C*_0_ is the eDNA concentration at the reference point, *v* the water velocity and *τ* the decay time (Altermatt et al., 2023; Sansom and Sassoubre, 2017). By assuming typical values such as *v* = 1 ms^−1^ and *τ* = 4 h (i.e., a half life of 2.77 h), we would be able to measure only about 50% of the concentration that we would measure 10 km upstream, and 3% of the concentration that we would measure 50 km upstream. Importantly, this estimate considers a channel with no lateral inputs; if lateral inputs with water not containing the target eDNA were considered, these percentages would inevitably decrease. As a rule of thumb, sites with a pairwise along-stream distance higher than 50 km should be considered as non-nested samples and thus unsuitable for eDITH’s spatial inference.

Finally, the model assumes that eDNA is well mixed in the water column, and/or that samples at a site adequately span the river cross section: for instance, sampling should be performed at both banks and at the centre for large rivers, while in small rivers sampling at a single bank might be sufficient, by assuming that eDNA is sufficiently well mixed. Analogous care should be taken in the choice of a representative sampling volume to be filtered (Altermatt et al., 2023).

#### Measurement errors

The eDITH model assumes that the sampling procedure applied is adequate with respect to sampling intensity (e.g., water volume sampled or sequencing depth; Altermatt et al. 2023; Bruce et al. 2021), and that subsequent laboratory procedures (e.g., DNA extraction, PCR, sequencing) do not introduce a systematic bias in DNA concentrations across samples. When using eDITH to model metabarcoding data, the expected read count at a given site for a given taxon is assumed to be proportional to the underlying DNA concentration. Replicated eDNA measures at the same site and time point are treated as independent measures. No distinction is made between physical (i.e., different water samples) or laboratory (different PCR runs) replicates.

#### What taxa can be modelled?

In principle, eDITH is suitable for aquatic eukaryotes (including invertebrates, vertebrates and plants) shedding their DNA into streamwater. Nonetheless, eDITH can also be extended for microorganisms sampled in their entirety (e.g., free floating bacteria), in which case their upstream sources can be considered as biofilm colonies, and the decay time does not refer to DNA molecules but rather to the bacteria lifetime. The eDITH model does not distinguish whether the eDNA data is referred to an assigned taxon (say, at a species, genus or family level) or an unassigned cluster such as OTUs or ASVs. In either case, the aforementioned assumptions on production and decay rates (see *DNA production and decay rates*) must hold. In principle, it is also possible to estimate the spatial distribution of terrestrial taxa (Lyet et al., 2021; Zong et al., 2023; Zhang et al., 2023), provided that the assumption of proportionality between DNA production rate and taxon density holds. In this case, the spatial unit where predictions are performed is the subcatchment, i.e., the portion of land that directly drains towards the associated reach.

#### River network model

The eDITH model can be applied to any river network discretized into reaches, that is, segments of the river not interrupted by confluences, which are treated as smallest spatial units. Each reach is considered as a network node, and the ensemble of all reaches covers the entire river network. It is assumed that reaches have no internal variability (e.g., the exact coordinates of a sampling site do not matter, provided that the sampling site is associated to the same reach). The along-stream distance between two consecutive confluences can be covered by a sequence of distinguished reaches, to allow for a finer discretization of the river network. Number and maximum length of reaches can be tuned via function `aggregate river` of the `rivnet` package (Carraro, 2023). See also Carraro et al. (2020a) and the documentation of the `OCNet` package for details on aggregation of a river network into reaches.

#### Lakes and braided channels

The eDITH model is designed for river networks not containing lakes, reservoirs, or systems where water is transferred against gravity and across drainage boundaries, e.g. through artificial channels and pumps. Dynamics of eDNA in lakes are still largely unclear, but the very large residence times of water particles in lakes (often months to years) compared to those of a river reach imply that all dissolved DNA entering a lake either degrades or is retained by substrate particles in the sediments. The river networks produced by `rivnet` do not admit bifurcations in the downstream direction (e.g., braided or artificial channels creating loops in the river network). Hence, the different braids or channels cannot be treated as independent reaches in eDITH. In this case, eDITH could still be applied by considering a single channel as a conceptual equivalent of the multiple real braids or channels.

### 2.2 The governing equations

The main equation of eDITH (Carraro et al., 2018, 2020b) results from a mass balance of eDNA across a river’s cross-section:

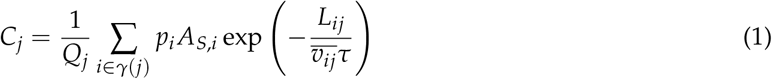

where *C*_*j*_ is the DNA concentration at a sampling site *j* (strictly speaking, *j* is the reach where a sampling site is located); *Q*_*j*_ is the water discharge in *j*; *γ*(*j*) identifies the set of reaches upstream of *j*; *p*_*i*_ is the DNA production rate at an upstream reach *i*; *A*_*S,i*_ is the source area in *i* (i.e., extent of the node; considering an aquatic taxon and a river reach, this could be equal to the product of the reach length and width); *L*_*ij*_ is the length of the along-stream path joining *i* to *j*; 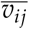 is the average water velocity along such path; *τ* is a characteristic decay time for DNA in stream water.

Assuming that river hydromorphology is known (and thus lengths, areas and discharges), Eq. (1) links DNA concentrations to the unknown parameters *τ* and **p** = (*p*_1_, …, *p*_*N*_), where *N* is the total number of reaches. While the former is linked to DNA behaviour in streamwater and could in principle be measured, or at least its value be constrained, the estimation of the latter is the actual goal of eDITH. Contrasting observed and modelled DNA concentrations thus enables the estimation of maps of **p** across all *N* reaches constituting the river network, and hence of relative taxon density, given the initial assumption.

It is often convenient, both from a modelling and interpretation viewpoint, to express the DNA production rate *p*_*i*_ via a Poisson generalized linear model as a function of environmental covariates, possibly related to the spatial patterns of the investigated taxon:

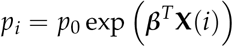

where **X**(*i*) is a vector of covariates evaluated at reach *i*; ***β*** a vector of covariate effect sizes; and *p*_0_ a baseline production rate, i.e., the DNA production rate at a site where all covariates are at a null level. This reduces the number of unknowns from *N* (length of **p**) to the number of selected covariates (length of ***β***) plus one (*p*_0_).

Importantly, if the eDNA data are expressed as read counts, Eq. (1) remains applicable. In this case, *C*_*j*_ would play the role of the expected read number at site *j* (proportional to the underlying DNA concentration, as previously hypothesized). Consequently, **p** would represent the DNA production rates multiplied by such constant of proportionality. For the sake of generality, in `eDITH` *C*_*j*_’s are termed “eDNA values”, which can either represent concentrations or read numbers, depending on the type of data used.

### 2.3 Estimating detection probability

Once model parameters are estimated (see *Model fitting and output analysis*), DNA production rates **p** can be transformed into corresponding detection probabilities **p**_**D**_ (vector of size *N*, i.e., one value per reach). This is done by exploiting the assumption on the probability distribution used to model measurement errors, and hence formulate the likelihood. In particular, for each reach *j*, we calculate the expected eDNA value 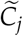 (concentration or read count) if that reach were detached from the river network, that is, in the absence of upstream inputs, with the water discharge in the reach being equal to the locally produced discharge (calculated as the actual discharge *Q*_*j*_ minus the sum of discharges of the upstream reaches that are directly connected to *j*). This is then transformed into a detection probability *p*_*D,j*_, calculated as the probability of observing a non-null eDNA value under the assumed error probability distribution and the expected value 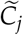. The so-obtained detection probability maps can be further transformed into presence-absence maps and used to assess biodiversity patterns (see e.g. Carraro et al. 2023).

## 3 The package structure and usability

The `eDITH` package consists of three main functions (Fig. 2): `run eDITH single`, which executes eDITH for a given set of parameters; `run eDITH optim`, which fits eDITH on provided data via non-linear optimization; and `run eDITH BT`, which fits eDITH following a Bayesian approach. In this latter case, functions `eval posterior eDITH` and `posterior pred sim eDITH` can be used for further output analysis (see *Model fitting and output analysis*).

**Figure 2:**
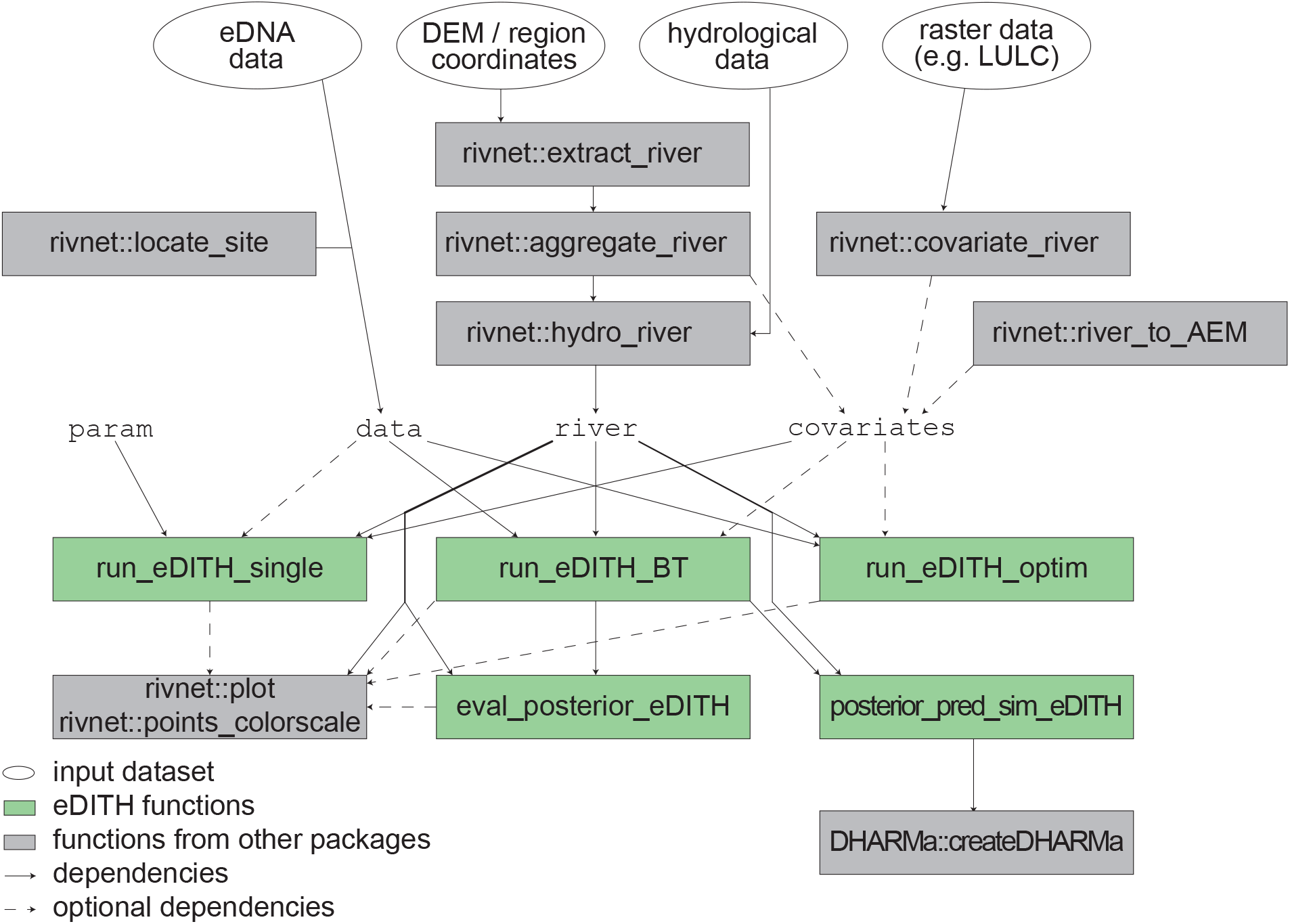
Workflow of the eDITH package and overview of its functions. Solid and dashed lines indicate mandatory and optional dependencies, respectively; for instance, covariates can be constructed by using at least one between aggregate river, covariate river, river to AEM from rivnet; covariates is a mandatory argument for run eDITH single but optional for run eDITH BT and run eDITH optim.

### 3.1 Required data

Data must be provided in the `data` field of `run eDITH BT` and `run eDITH optim` (and optionally for `runeDITH single`) as a data frame with components `values`, i.e., eDNA values measured for a given taxon and a given time point, and `ID`, i.e., identifiers of the network nodes at the aggregated (AG) level (i.e., reaches; see Carraro et al. 2020a) where the sampling sites are located. To identify these network nodes, function `locate site` from `rivnet` can be used.

The river network must be provided as a `river` object, obtained via the `rivnet` package. For this purpose, function `extract river` (Fig. 2) extracts a river network from either user-provided or opensource elevation data. In the latter case, it is sufficient to provide coordinates of the extent of the region of interest and location of the watershed outlet in a projected coordinate system, as well as the desired resolution of the digital elevation model (argument `z`). It is fundamental that the river be aggregated into reaches (via `aggregate river`) and that it contain hydrological data, such that discharges and water velocities could be used as input in eDITH. This is done via the `hydro river` function of `rivnet`. In its simplest setting, a single value of discharge (or depth) and width, not necessarily at the same reach, are required in order to extrapolate hydraulic variables across the whole network; ideally (and in more advanced settings) discharge data from multiple gauging stations can be used as input in `hydro river`. The exact method with which `hydro river` extrapolates hydrological variables depends on the number and type of data provided.

Optionally, covariates can be passed to `run eDITH BT` and `run eDITH optim` as a data frame. Function `covariate river` of `rivnet` allows computing covariates from raster maps and a `river` object. If covariates are not provided, asymmetric eigenvector maps (AEMs) are calculated on the river network and used as covariates. AEMs (Blanchet et al., 2008) are mutually orthogonal spatial variables obtained by a spatial filtering technique that considers space in an asymmetric way, and are thus suitable to model species distributions in river networks. It is of course possible to combine user-provided covariates and AEMs (Fig. 3). Covariates are a mandatory argument for `run eDITH single`, together with a vector of parameters `param`.

**Figure 3:**
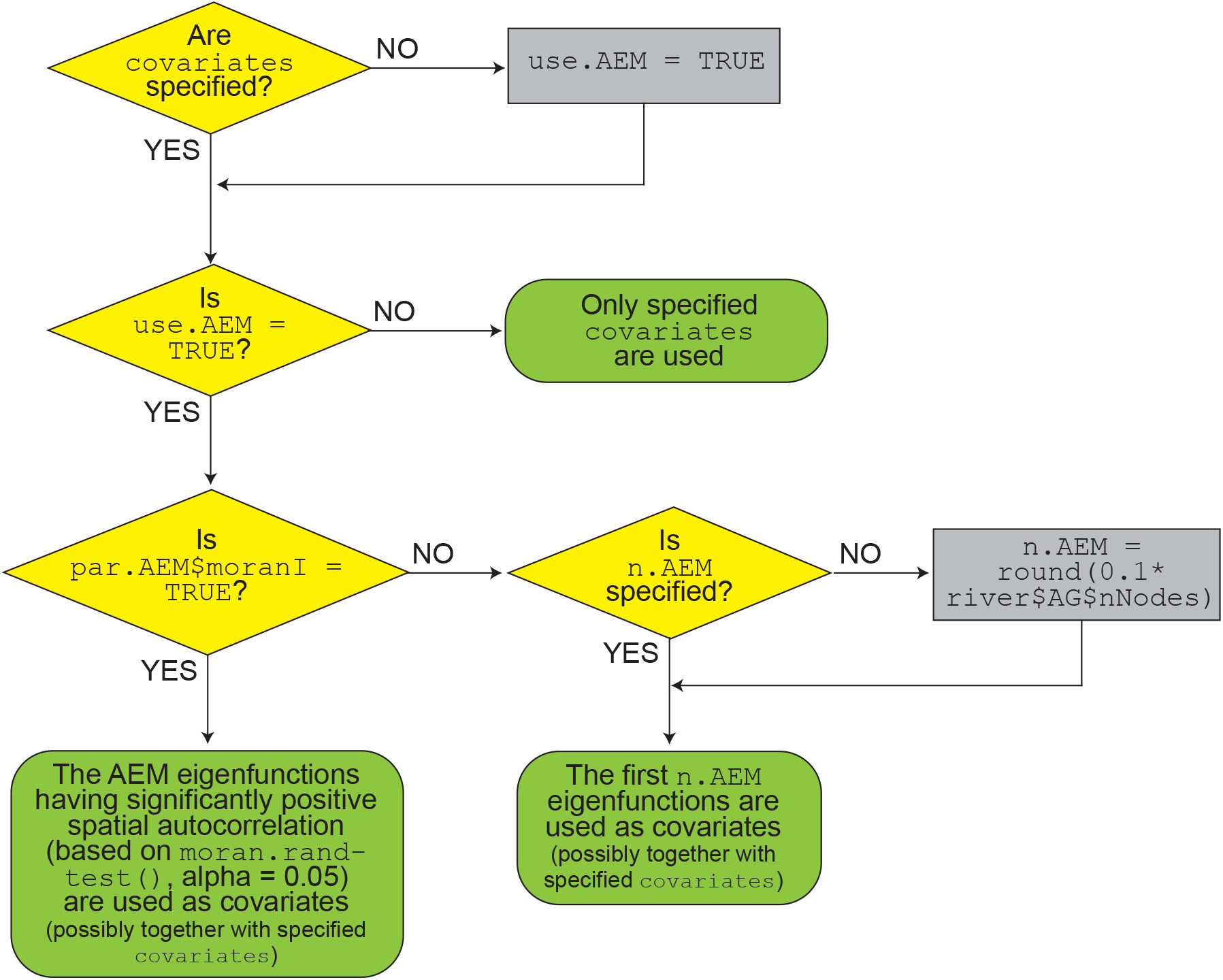
Flowchart for the choice of covariates used to fit eDITH.

### 3.2 Model fitting and output analysis

Function `run eDITH BT` makes use of a Bayesian sampler and thus interfaces with the `BayesianTools` Rpackage (Hartig et al., 2019), which provides a wide choice of Bayesian methods, with the default sampler in `eDTIH` being DREAM_ZS_ (Vrugt et al., 2009). Specifically, Bayesian methods allow sampling the posterior distribution of the model parameters; in `eDTIH`, both components constituting the posterior distribution, i.e., the likelihood function and the prior distribution, can be specified by the user.

Four probability distributions are implemented in `eDITH` to model errors between observed and modelled data and thus calculate the likelihood: the normal (`ll.type = “norm”`) and log-normal (`ll.type = “lnorm”`) distributions are suitable for DNA concentration data, such as measured via qPCR; conversely, the geometric (`ll.type = “geom”`) and negative binomial (`ll.type = “nbinom”`) distributions are suitable for read count data, i.e., produced by metabarcoding. Importantly, the user is expected to set option `ll.type` according to the nature of the `data` provided in `run eDITH BT` or `run eDITH optim`. The optional argument `no.det` can be used to produce a zero-inflated probability distribution. This is mandatory if `ll.type = “lnorm”`, as the log-normal distribution does not admit zeros (see Carraro et al. 2018).

Default model parameters and their respective prior distributions are as follows. Firstly, `tau` represents decay time *τ* in hours, and its default prior is a log-normal distribution with mean of 5 h and median of 4 h. Secondly, `log p0` is the decimal logarithm of the baseline production rate *p*_0_; with a default uniform prior bound between −20 and 0. The units of *p*_0_ are equal to the units of the inputted `data`$`values` multiplied by a length unit and divided by a time unit. For instance, if `data`$`values` contains DNA concentration data in mol m^−3^ and the `river` object contains discharges in m^3^s^−1^ and areas in m^2^, *p*_0_ is expressed in mol m^−2^ s^−1^ (i.e., amount of DNA shed per unit area and unit time). In the case of read count data, the unit for *p*_0_ is ms^−1^; this is because, in this case, *p*_0_ embeds a constant that transforms expected DNA concentration predicted by the model into expected read number. Finally, `beta X` is the effect size of covariate `X`. Default priors for the effect size parameters are normal distributions with null mean and unit standard deviation. By default, all covariates are Z-normalized.

Additional parameters can be present depending on the likelihood definition. When a normal or lognormal error distribution is chosen for the likelihood, parameter `sigma` identifies the standard deviation of the error. When a negative binomial error distribution is used, `omega` represents the overdispersion parameter, that is the ratio between variance and mean (Lindén and Mäntyniemi, 2011). No additional error parameter is needed when a geometric error distribution is used. Finally, `Cstar` is a further parameter added when a zero-inflated probability distribution is chosen (i.e., when `no.det = TRUE`). This represents a characteristic eDNA value *C*^*^ below which the probability of a non-detection becomes relevant (Carraro et al., 2018). Specifically, the non-detection probability at a given site *j* is calculated as exp(−*C*_*j*_/*C*^*^), where *C*_*j*_ is the modelled eDNA value in *j*. Increased *C*^*^ increases the zero-inflation of the probability distribution of observed values, all else being equal.

A faster alternative to Bayesian inference is constituted by `run eDITH optim`, which seeks to maximize the posterior distribution via non-linear optimization; to achieve this, function `optim` from base R is called. By default, several optimization attempts are made starting from randomly chosen initial parameter sets, in order to diminish the probability that the optimization algorithm remains trapped in a local maximum. The choice of maximizing the posterior distribution rather than the likelihood enables restricting the parameter set to physically meaningful ranges (in particular, *τ* must be positive).

Output from `run eDITH BT` and `run eDITH optim` is a list containing best-fit parameters (maximum-aposteriori estimates for `run eDITH BT`) and the respective estimates of eDNA values **C**, production rates **p** and detection probabilities **p**_**D**_. To help further analysis, output from the respective fitting functions (`runMCMC` from `BayesianTools`, `optim`) is also exported. When the Bayesian approach is used, functions `eval posterior eDITH` and `posterior pred sim eDITH` allow calculating relevant quantiles from the parameters’ posterior distribution, and running posterior predictive simulations, respectively. Specifically, the latter can be used for diagnostics purposes such as assessing scaled (quantile) residuals via the `DHARMa` package. Finally, estimated **C, p** and **p**_**D**_ can be visualized across the river network via the `plot` method and the `points colorscale` function of `rivnet` (Fig. 2).

## 4 Case studies

We illustrate `eDITH`’s functionalities by applying it to two published case studies of spatially replicated fish eDNA metabarcoding datasets across two catchments, the Lower East Sydenham river (Canada, 1623 km^2^, 30 sampling sites; Balasingham et al. 2018) and the Koide river (Japan, 39 km^2^, 11 sampling sites; Sakata et al. 2021). A detailed explanation and a step-by-step protocol of the model implementation is provided in an R Markdown file (see *Data availability statement*). In brief, following the workflow of Fig. 2, we delineated the two watersheds using open source elevation data in `rivnet`. We used the first 10 AEMs as covariates, and fitted eDITH via `run eDITH BT` by using a negative binomial error distribution and default settings for the other arguments. For both river systems, we restricted our analysis to species only found in at least 5 sampling sites. For each modelled species, we calculated the median of the posterior distribution of detection probabilities via `eval posterior eDITH` (see a snippet of results in Fig. 4) and implemented a posterior predictive check via `posterior pred sim eDITH` and `createDHARMa`. Finally, we determined species’ presence/absence with a threshold of 0.5 on the median posterior detection probability, and produced maps of species richness that we compared with the observed species richness (Fig. 4).

**Figure 4:**
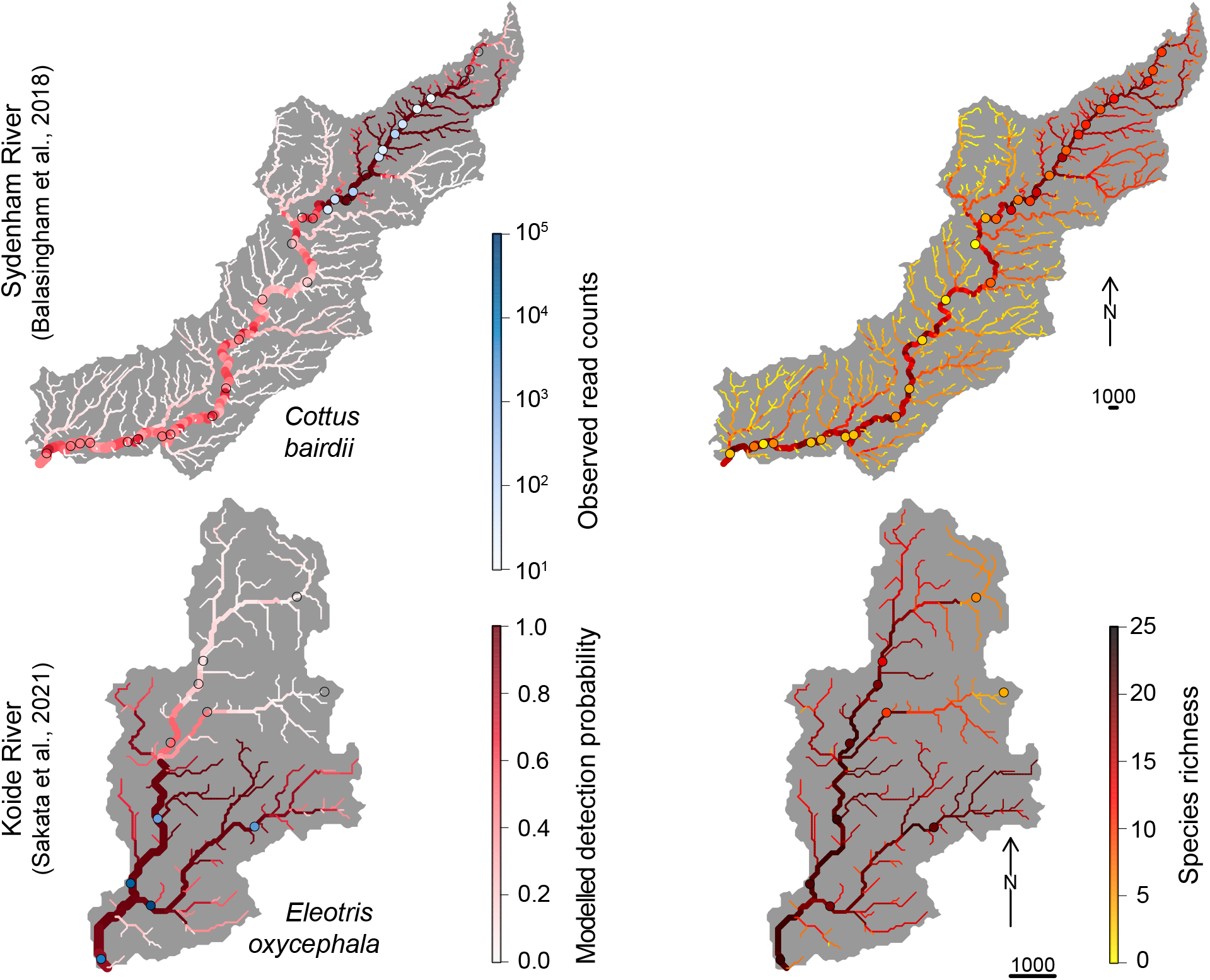
Results from the application of eDITH to the datasets of (Balasingham et al., 2018) (top row) and (Sakata et al., 2021) (bottom row). Left column: modelled detection probability (median of the posterior distribution) for a selected fish species shown along the river network via rivnet’s plot; observed read counts for the same species displayed as points via rivnet’s points colorscale. Transparent colouring indicates non-detection. Right column: modelled species richness shown along the river network; observed species richness displayed as points.

## 5 Conclusions

Freshwater ecosystems are among the most biodiverse but also most impacted by humans (Dudgeon, 2019), and are thus subject to intense monitoring including biodiversity and ecological indices assessments. Novel eDNA-based approaches, especially when coupled with hydrology-based models, offer unprecedented opportunities to advance freshwater biodiversity assessments. In particular, the eDITH model exploits spatially replicated eDNA measurements within a river to infer the spatial distribution of any taxon of interest. Its underlying concept is that eDNA particles are advected by streamflow, and hence an eDNA sample not only represents the sampling location, but provides information on a certain upstream area. By exploiting multiple, spatially distributed sampling sites, and considering DNA transport and concurrent decay, eDITH can disentangle the various sources of DNA shedding. Therefore, eDITH complements point-wise eDNA measurements by projecting taxon distributions into space-filling catchment maps. To allow a broad application of eDITH by a wider community of scientists and practitioners, we presented the `eDITH` R-package, which simplifies the implementation of all necessary steps for the use of eDITH, from watershed delineation to model fitting and visualization. We expect this tool to strongly contribute to a more precise and upscaled assessment of freshwater ecosystems, applicable across all riverine systems globally with minimal prior information, and thus also accessible for undermonitored regions.

## Data availability statement

The `eDITH` package is available on CRAN: https://cran.r-project.org/web/packages/eDITH/index.html. A commented script illustrating the case study examples is available on Github: https://github.com/lucarraro/test_eDITH.

## Acknowledgements

Funding is from the Swiss National Science Foundation (Ambizione grant PZ00P2 202010 to L.C. and grant 31003A 173074 to F.A.) and the University of Zurich Research Priority Programme URPP GCB (to F.A.).

